# Selective Detection of a Key Region in Chemotaxis Signaling Protein Complexes by Solid-State NMR

**DOI:** 10.1101/2025.05.15.654355

**Authors:** Jessica J. Allen, Songlin Wang, Chad M. Rienstra, Lynmarie K. Thompson

**Affiliations:** Department of Chemistry, University of Massachusetts Amherst, Amherst, Massachusetts, 01003; National Magnetic Resonance Facility at Madison (NMRFAM), University of Wisconsin-Madison, Madison, Wisconsin, 53706; Department of Chemistry, University of Wisconsin-Madison, Madison, Wisconsin, 53706; Department of Biochemistry, University of Wisconsin-Madison, Madison, Wisconsin, 53706; Morgridge Institute for Research, Madison, Wisconsin, 53706

## Abstract

Understanding how large protein complexes function requires tools that can resolve both structure and dynamics in their native assembly states. Here, we apply solid-state NMR (SSNMR) to selectively detect rigid regions of a receptor protein fragment in the context of the >500 kDa chemoreceptor signaling complex found in chemotactic bacteria. These complexes assemble into hexagonal arrays that network multiple active units together. The cytoplasmic fragment of the *E. coli* aspartate chemoreceptor exhibits dynamics on multiple timescales across different regions of the protein, and these dynamics differ between signaling states. We apply ^13^C-^15^N dipolar coupling-based SSNMR experiments to selectively probe the rigid portion (motions slower than millisecond timescale) of this protein, in the context of the full array structure. We optimized assembly methods to form native-like, homogeneous complexes capable of maintaining activity and sample integrity during extended NMR experiments with low electric-field NMR probe designs. A subset of the protein, approximately 100 residues at the membrane-distal tip of the chemoreceptor where it interacts with its associated kinase, was detected and identified as the most rigid region. Chemical shift changes for many residues in this rigid region were observed between NMR spectra of the kinase-on and kinase-off signaling states. This suggests conformational changes occur at the chemoreceptor tip during signaling, which have not been observed in previous studies of this system. These findings demonstrate a dynamics-based NMR spectral editing approach to selectively examine a key region of a large signaling protein within its macromolecular assembly.

**Statement of Significance:** Large protein complexes are challenging to study, but understanding their mechanism is vital to gaining insight into many important biological processes. In this work we demonstrate the use of solid-state NMR to study such a system, bacterial chemotaxis receptor signaling complexes. We identified an *in vitro* assembly method that creates reproducible and durable complexes that retain activity for weeks of data collection. Selective NMR detection of a key region of one protein reveals significant differences between signaling states, indicating there are changes in conformation and dynamics that were not previously seen by cryo-electron tomography and crystallography of this system. This highlights the potential of solid-state NMR as a key tool in mechanistic studies of multi-protein complexes.

## Introduction

Understanding the full mechanistic picture for large protein assemblies requires insights into both the dynamics and conformational landscapes of their component proteins within functional complexes. X-ray crystallography and cryo-electron microscopy (cryo-EM) are essential tools for determining high resolution structures of proteins and protein complexes, but they often struggle to capture dynamic or disordered regions, which are critical for biological function. Nuclear magnetic resonance (NMR) is a powerful technique capable of measuring both structure and dynamics of proteins with atomic resolution. Recent advances, including novel pulse sequences, labeling strategies, and fast magic angle spinning (MAS), have allowed structure and dynamics of HSP90-Tau complex (1), ClpB-DnaK complex (2) and the HIV-1 capsid (3) to be studied using solution and solid-state NMR (SSNMR). Magic-angle spinning (MAS) SSNMR is especially useful for large protein assemblies or aggregates with slow tumbling. One example is an SSNMR study of the type III secretion system, a needle structure that extends from the membrane of pathogenic bacteria into the extracellular space, and consists of multiple copies of an ∼80-residue subunit protein(4). The homogeneity and rigidity of the needle structure allowed for near-complete assignment of dipolar coupling-based 2D ^15^N/^13^C correlation experiments, providing insights into secondary structures and the conformation of the subunit protein within the needle(5). Studies like this highlight how SSNMR has expanded the range of biological structures that can be analyzed at the residue level to reveal both structural and dynamic information.

A major challenge in NMR studies of large proteins and protein assemblies is spectral crowding due to the large number of residues in similar secondary structures. This has led to the advent of various isotopic labeling techniques, including specific, reverse, and sparse labeling, to reduce the number of labeled sites and thereby simplify the spectra (6–9). In addition to these labeling strategies, another complementary approach for selectively detecting certain regions of uniformly labeled protein takes advantage of different timescales in molecular motion within a single protein. Scalar-based NMR experiments, such as INEPT, detect signals from regions with motions on the nanosecond or faster timescale, while dipolar-coupling based NMR experiments, like cross polarization (CP), identify regions that are rigid on the nanosecond timescale (10–13). This can relieve spectral congestion by separating the signals into distinct spectra for dynamic and rigid regions. For instance, our group has applied this approach for selective detection of the most dynamic portion of the aspartate chemoreceptor in large signaling complexes, enabling amino acid type assignments and revealing changes in the number of highly dynamic residues between signaling states (14, 15).

*E. coli* chemoreceptor signaling complexes provide the sensory input for bacterial chemotaxis and are well-studied systems for investigations of transmembrane signaling mechanisms. The chemoreceptor is a membrane-spanning receptor that binds ligands in the environment and transmits signals across the membrane to control a histidine kinase, CheA (Figure 1A). The core signaling unit, the smallest assembly required for activity of these complexes, is composed of two trimer-of-dimers of receptors, a homodimer of CheA, and two monomers of the coupling protein CheW (Fig. 1B). The signaling state of the complex is determined by ligand binding to the receptor periplasmic domain and methylation of four specific glutamate residues in the receptor cytoplasmic domain. Attractant binding and demethylation of the chemoreceptor result in the inhibition of the kinase. However, the overall mechanism of how the chemoreceptor controls the kinase activity remains unclear.

**Figure 1.**
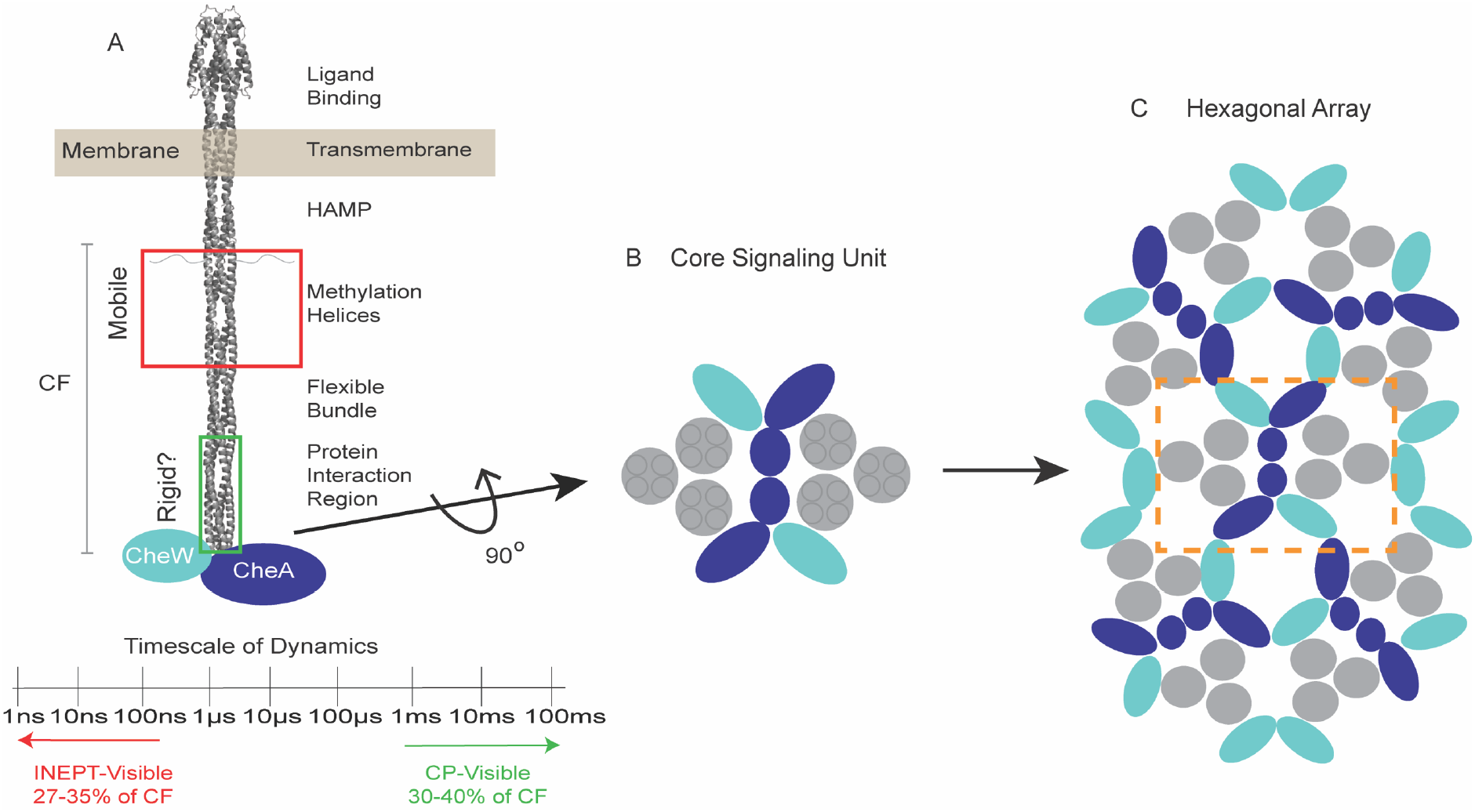
Overview of structure and dynamics of chemoreceptor signaling complexes and arrays. (A) Overview of domains of a chemoreceptor dimer (gray). The region identified to be highly dynamic (INEPT-Visible) by previous SSNMR studies is in the red box, and the region predicted to be the most rigid, and thus observable in CP NMR spectra, is in the green box. The cytoplasmic fragment (CF) of the chemoreceptor includes the methylation helices, flexible bundle, protein interaction region (PIR) and C-terminal tails. (B) Bottom-up view of a core signaling unit (CSU) comprised of two chemoreceptor trimer-of-dimers (gray), one CheA homodimer (blue) and two CheW (cyan). (C) Bottom-up view of hexagonal array that forms at the poles of chemotactic bacteria. Neighboring CSUs are connected through CheW-CheA and CheW-only rings.

Core signaling units interact via rings of the CheA and CheW proteins to further assemble into large hexagonal arrays with multiple core units (Fig 1C) (16). Cryo-electron tomography (Cryo-ET) combined with X-ray crystallography and molecular dynamics simulations have produced structural models of these complexes. These models are limited to a resolution of 8.4 Å and do not detect any differences between the signaling states in the protein interaction region where the chemoreceptor forms contacts with CheA and CheW (17). Resolving these interactions at atomic level and understanding how they change between signaling states is essential for uncovering the mechanism of signaling.

Measurement of differences in structure and dynamics between the kinase-on and kinase-off signaling states requires homogeneous, functional signaling complexes. Several established protocols are available to prepare native-like signaling arrays *in vitro*, which involve combining an aspartate receptor cytoplasmic fragment (CF) with CheA, CheW and agents that mediate spontaneous assembly of the complexes (18–21). The methylation helices of the CF contain the glutamic acid residues that undergo methylation and demethylation to regulate kinase activity as part of adaptation during chemotaxis. CF with 4 glutamate residues (CF4E) inhibits the kinase, while a methylated-mimic of CF with glutamate replaced by glutamine (CF4Q) activates the kinase. Thus, assemblies made with CF4E or CF4Q represent kinase-off and kinase-on states, respectively (18).

In this work, we demonstrate the selective detection and identification of the most rigid region of the CF in functional complexes, which is a key region for understanding the signaling mechanism. Previous SSNMR INEPT experiments in our lab on CF incorporated into these complexes have shown that the methylation helices and C-terminal tail have nanosecond or faster motions (14, 15). Dipolar coupling-based NMR experiments such as ^1^H to ^13^C CP experiments provide a means to observe the rigid fraction of the protein. CP buildup rates are comparable for CF in frozen samples— assumed to represent 100% rigid conditions—and unfrozen samples, but intensities reach only 25-40% (depending on signaling state) in unfrozen samples (15, 22). This suggests that only some region(s) of the CF are rigid enough to be observed in CP spectra. As summarized in Figure 1A, ∼1/3 of the CF residues are observable in INEPT spectra and ∼1/3 are observable in CP experiments. Furthermore, ^15^N-^13^C REDOR experiments have shown that only a fraction of the CP-observable residues have rigid-limit CN dipolar couplings (22). This predicts that dipolar coupling-based experiments, such as ^15^N/^13^C_α_ correlation (NCA) and ^15^N/^13^CO correlation (NCO) experiments, will detect signals from a subset of the protein, roughly 80 of its 310 residues. ^1^H/^2^H exchange (HDX) mass spectrometry by our lab has shown that the protein interaction region (PIR) of the chemoreceptor, which interacts with CheA and CheW, has the slowest deuterium uptake (23). This suggests that the 57-residue PIR is part of the most structured and least dynamic portion of the protein. In this work, we show that ^13^C-^15^N dipolar coupling-based experiments selectively detect signals from part of the CF and identify this region as the PIR as well as an adjacent portion of the flexible bundle. Comparison of these experiments done on kinase-on and kinase-off states of the functional chemoreceptor signaling complexes reveal numerous chemical shift changes, widespread through the sequence, suggesting that many residues have different interactions and undergo conformational changes between signaling states.

## Materials and Methods

### Protein Expression

The pTEV-cheA (kan^r^) and pTEV-cheW (kan^r^) were expressed in BL21(DE3) *E. coli* cells for production of unlabeled CheA and CheW (14). The plasmid pHTCF4Q (amp^r^) encodes the methylated-mimic of the cytoplasmic fragment (CF) with glutamine at all 4 methylation sites, while pHTCF4E (amp^r^) encodes the unmethylated CF with glutamate at those sites (18). These plasmids were expressed in BL21(DE3) cells in minimal media containing ^13^C-glucose and ^15^N-ammonium sulfate for uniform isotopic labelling (14). All four proteins contain N-terminal His tags and were purified using nickel affinity chromatography; the TEV cleavable His-Tags were removed from the CheA and CheW. All protein expression and purification were performed as previously reported (21).

### Pellet Stability Studies

PEG-mediated assemblies were made by combining, in order, low salt kinase buffer (30 mM K_x_H_x_PO_4_ and 15 mM KCl, pH 7.5), 12 µM CheA, 20 µM CheW, 1 mM PMSF, 7.5% w/v PEG8000 and 50 µM CF. Vesicle assemblies were made by combining, in order, kinase buffer (50 mM K_x_H_x_PO_4_ and 75 mM KCl, pH 7.5), 12 µM CheA, 24 µM CheW, 1 mM PMSF, 725 µM lipids and 30 µM CF (14). For both assembly methods CF4Q was used to assemble a kinase-on state and CF4E was used to assemble a kinase-off state. 500 µL of each assembly was made for each stability study. Samples were incubated overnight at 25°C to allow for full array assembly.

Kinase, methylation and sedimentation assays were performed as previously described to ensure assemblies have formed properly (21). Kinase activity for PEG8000-mediated and vesicle assemblies ranged between 8-14 ATP consumed per total CheA (s^−1^) when measured after about 24 hours of assembly formation. Stoichiometries of the proteins from sedimentation assays for both assembly types were approximately 6CF:2CheW:1CheA, as observed for *in vivo* arrays. The methylation assay was performed by combining 6 µM CheR and 10 µM SAM with kinase-off assemblies to a final CF4E concentration of 39.5 µM, and then incubated at 25°C. Aliquots were removed after 10 minutes and 4 hours and the methylation reaction was quenched by adding an equal volume of 2X SDS gel sample buffer. Comparison samples consisted of complexes incubated without CheR and SAM. Samples were loaded on an 12.5% SDS PAGE gel for electrophoresis at 120V for about 80 minutes, followed by staining with Coomassie blue. The resolved bands of the methylated and unmethylated CF were quantified using ImageJ analysis. After 4 hours of methylation about 60% of CF4E is methylated.

A small aliquot of each assembly was kept in solution to use as a control, and multiple 50µL aliquots were ultracentrifuged at 128,000*g* for 1 hour. After ultracentrifugation, each supernatant was removed from the pellet, and both were stored separately in sealed Eppendorf tubes at 4°C. Activity measurements were performed after a range of incubation times. Pellets were resuspended, with their supernatant, the same day as the kinase or methylation assay. After the supernatant was added back to the pellet, the sample was incubated at room temperature for about an hour and then vigorously mixed, by pipetting up and down, to ensure the full resuspension of the pellet. Kinase activity assays were performed at each time point on kinase-on (CF4Q) resuspended pellets and compared to control samples of the same assembly kept in solution. Methylation assays were performed at each time point on kinase-off (CF4E) resuspended pellets and compared to control samples of the same assembly kept in solution.

### SSNMR Sample Preparation

SSNMR samples were prepared as described above for PEG-mediated assemblies but with 50 µM U-^13^C,^15^N CF (CF4Q for the kinase-on state or CF4E for the kinase-off state) and to a final volume of 4 mL. This created sufficient pellet material to fill two 1.6 mm SSNMR rotors. Kinase and methylation activity measurements, as well as sedimentation assays, were performed to ensure assemblies formed properly, and results were consistent with those reported above. Ultracentrifugation was used to pellet the assemblies for rotor packing. Samples were spun at 128,000*g* for 2 hours to form a gel-like pellet. The pellets were transferred from the ultracentrifuge tubes by scraping and centrifuging into a rotor packing device specifically designed for the rotor size, then packed into either 1.6 mm (8 µL volume) or 2.5 mm (18 µL volume) rotors following a standard protocol (24, 25). Amount of sample packed was determined from the change in weight of the rotor before and after packing, with a resulting estimate of about 70 or 170 nmoles of CF packed in the 1.6 mm or 2.5 mm rotors, respectively.

### NMR Data Collection

All spectra for samples packed in 1.6 mm rotors were collected using a Bruker Avance III HD 900 MHz spectrometer (21.1T) with a Black Fox (Tallahassee, FL) probe with PhoenixNMR (Loveland, CO) 1.6 mm spinning module. The probe was tuned to HCN mode with optimization of the ^13^C sensitivity (26). One spectrum of a sample packed in a 2.5 mm rotor (SI Figure 3) was collected using a Bruker NEO 900 MHz spectrometer (21.1T) with a Black Fox probe with PhoenixNMR 2.5 mm spinning module. The sample spinning rates were 20 kHz for all experiments. The variable temperature (VT) calibration was completed using methanol (27). The VT gas temperature was set to -5°C, resulting in an actual sample temperature of 4°C. It is notable that both Black Fox probes utilize low electric field resonators for the ^1^H channel operation, reducing the transient heating of the samples during long decoupling pulses (28, 29). All spectra were referenced to DSS, using adamantane as a secondary external standard. The left ^13^C signal of adamantane was referenced to 40.48 ppm (30).

For the 2D ^13^C/^13^C correlation experiment (CC), ^13^C polarization was prepared using adiabatic CP with a downward tangential ramp pulse applied on the ^1^H channel. The CP contact time was 0.6 ms, with a ^13^C radio frequency (RF) amplitude of 90 kHz and an average ^1^H RF amplitude of 69 kHz. Polarization transfer between ^13^C nuclei was achieved using dipolar-assisted rotational resonance (DARR) mixing of 50.0 ms, with a ^1^H RF amplitude of 20 kHz (31). The t_1_ acquisition time was 6.4 ms, with a 12.5 µs increment over 512 complex points. The t_2_ acquisition time was 15.4 ms, with a 5 µs dwell time for 1536 complex points. During the acquisition, SPINAL-64 ^1^H decoupling at 100 kHz was applied. The total number of scans is 32, and the recycle delay was 1.1 s. The 2D spectrum was acquired using a 25% non-uniform sampling (NUS) schedule; total experimental time was 10.8 hours (32).

The 2D ^15^N/^13^C_α_ correlation experiment (NCA) was performed with the ^13^C carrier frequency set at 55 ppm using SPECIFIC CP (33). The ^15^N polarization was prepared via an adiabatic CP with a downward tangential ramp pulse applied on the ^1^H channel, with a contact time of 1.4 ms, ^15^N RF amplitude of 30 kHz, and an average ^1^H RF amplitude of 69 kHz. ^15^N polarization was then transferred to ^13^C_α_ using CP with an upward tangential ramp on the ^13^C channel, with a contact time of 3.5 ms, ^15^N RF amplitude of 30 kHz, and an average ^13^C RF amplitude of 12 kHz. CW decoupling of ^1^H at 100 kHz was applied during this period. The t_1_ acquisition time was 12.8 ms with a 50 µs increment for 256 complex points, while a 5.1 µs ^13^C π-pulse was applied at the center of the t_1_ period to decouple ^1^*J*_*NC*_. The t_2_ acquisition time was 20.5 ms with a 5 µs dwell time for 2048 complex points. SPINAL-64 ^1^H decoupling at 100 kHz was applied during acquisition. The total number of scans is 64, and the recycle delay was 1.1 s. The 2D spectrum was acquired using a 25% NUS schedule; total experimental time was 10.5 hours.

The 2D ^15^N/^13^CO correlation experiment (NCO) was performed with the ^13^C carrier frequency set at 175 ppm. The ^15^N polarization was prepared via an adiabatic CP with a downward tangential ramp pulse applied on the ^1^H channel, with a contact time of 1.2 ms, ^15^N RF amplitude of 46 kHz, and an average ^1^H RF amplitude of 84 kHz. ^15^N polarization was then transferred to ^13^C_α_ using CP with an upward tangential ramp on the ^13^C channel, with a contact time of 7.0 ms, ^15^N RF amplitude of 30 kHz, and an average ^13^C RF amplitude of 52 kHz. CW decoupling of ^1^H at 100 kHz was applied during this period. The t_1_ acquisition time was 12.8 ms with a 100 µs increment for 128 complex points, while a 5.1 µs ^13^C π-pulse was applied at the center of the t_1_ period to decouple ^1^*J*_*NC*_. The t_2_ acquisition time was 15.4 ms with a 5 µs dwell time for 1536 complex points. SPINAL-64 ^1^H decoupling at 100 kHz was applied during acquisition (34). The total number of scans is 64, and the recycle delay was 1.1 s. The 2D spectrum was acquired using a 25% NUS schedule, total experimental time was 7.1 hours.

The 3D ^15^N/^13^C_α_/^13^CX correlation experiment (NCACX) was performed with the ^13^C carrier frequency set at 55 ppm. The ^1^H to ^15^N CP and the ^15^N to ^13^C_α_ CP transfer conditions were identical to the 2D NCA experiment as described above. ^13^C_α_ to ^13^CX polarization transfer was achieved using a 50.0 ms DARR mixing period with a ^1^H RF amplitude of 20 kHz. The t_1_ acquisition time was 9.6 ms with a 150 µs increment for 64 complex points, and a 5.1 µs ^13^C π-pulse was applied at the center of the t_1_ period to decouple *J*_*15N-13C*_. The t_2_ acquisition time was 6.4 ms with a 100 µs increment for 64 complex points. At the center of the t_2_ period, a 5.1 µs ^13^C hard π-pulse, a 275 µs ^13^C soft π-pulse with rSNOB shape at 55 ppm, and an 9.0 µs ^15^N π-pulse were applied to decouple the *J*_*13Cα-13CX*_ and *J*_*13Cα-15N*_ (35). The t_3_ acquisition time was 20.5 ms with a 5 µs dwell time for 2048 complex points. SPINAL-64 ^1^H decoupling at 100 kHz was applied during acquisition. The total number of scans is 88, and the recycle delay was 1.1 s. The 3D spectrum was acquired using a 25% NUS schedule, and the total experimental time was 120 hours.

The 3D ^15^N/^13^CO/^13^CX correlation experiment (NCOCX) was performed with the ^13^C carrier frequency set at 175 ppm. The ^1^H to ^15^N CP and the ^15^N to ^13^C_α_ CP transfer conditions were identical to the 2D NCO experiment as described above. ^13^CO to ^13^CX polarization transfer was achieved using a 50.0 ms DARR mixing period with a ^1^H RF amplitude of 20 kHz. The t_1_ acquisition time was 9.6 ms with a 150 µs increment for 64 complex points, and a 5.1 µs ^13^C π-pulse was applied at the center of the t_1_ period to decouple *J*_*15N-13C*_. The t_2_ acquisition time was 7.2 ms with a 150 µs increment for 48 complex points. At the center of the t_2_ period, a 5.1 µs ^13^C hard π-pulse, a 275 µs ^13^C soft π-pulse with rSNOB shape at 175 ppm, and an 9.0 µs ^15^N π-pulse were applied to decouple the *J*_*13CO-13CX*_ and *J*_*13CO-15N*_. The t_3_ acquisition time was 20.5 ms with a 5 µs dwell time for 2048 complex points. SPINAL-64 ^1^H decoupling at 100 kHz was applied during acquisition. The total number of scans is 176, and the recycle delay was 1.1 s. The 3D spectrum was acquired using a 25% NUS schedule, and the total experimental time was 180 hours.

The 3D CANCO experiment was performed with the ^13^C carrier frequency set to 175 ppm. ^13^C_α_ polarization was prepared via adiabatic CP using a downward tangential ramp pulse on the ^1^H channel, with the ^13^C frequency adjusted to 55 ppm for ^1^H-^13^C_α_transfer. The CP contact time was 0.6 ms, with a ^13^C RF amplitude of 90 kHz, and an average ^1^H RF amplitude of 69 kHz. Polarization was then transferred from ^13^C_α_ to ^15^N via CP using an upward tangential ramp on the ^13^C channel. The contact time was 6.0 ms, with an ^15^N RF amplitude of 30 kHz, and an average ^13^C RF amplitude of 12 kHz. CW ^1^H decoupling at 100 kHz was applied during this CP period. The ^15^N polarization was transferred to ^13^CO using CP with an upward tangential ramp on the ^13^C channel after setting the ^13^C frequency back to 175 ppm. The contact time was 5.0 ms, with a ^15^N RF amplitude of 30 kHz, and an average ^13^C RF amplitude of 52 kHz. CW ^1^H decoupling at 100 kHz was applied during this CP period. The t_1_ acquisition time was 6.4 ms with a 100 µs increment for 64 complex points. A 5.1 µs ^13^C hard π-pulse, a 275 µs ^13^C soft π-pulse with rSNOB shape at 55 ppm, and an 9.0 µs ^15^N π-pulse were applied at the center of the t_1_ period to decouple the *J*_*Cα-CX*_ and *J*_*13Cα-15N*_. The t_2_ acquisition time was 9.6 ms, with a 150 µs increment for 64 complex points. A 5.1 µs ^13^C π-pulse was applied at the center of the t_2_ period to decouple the *J*_*N-C*_. The t_3_ acquisition time was 20.5 ms, with a 5 µs dwell time for 2048 complex points. SPINAL-64 ^1^H decoupling at 100 kHz was used during acquisition. The total number of scans is 148, and the recycle delay was 1.1 s. The 3D spectrum was acquired using a 25% NUS schedule, and the total experimental time was 194 hours.

### NMR Data Analysis

The 2D and 3D datasets were initially processed using NMRPipe, followed by data analysis and signal assignment using POKY (36, 37). Spectral processing included zero filling in all dimensions, linear prediction in the indirect dimension for 2D spectra (256 points for NCA, 128 points for NCO and 512 points for CC), polynomial baseline correction and 45° sine-bell window functions. To ensure a direct comparison of the 2D NCA spectra of CF4Q and CF4E, identical acquisition and processing parameters were applied. SMILE was used for NUS reconstruction for all 3D experiments (38). For visualization, contour levels were set to 7 times the noise level. 2D NCA, NCO and CC DARR experiments, as well as 3D NCACX, NCOCX and CANCO experiments, were used for signal assignment. The assignment results are summarized in Table S1 and S2.

## Results

### Optimized Sample Conditions for SSNMR of Functional Signaling Complexes

NMR resonance assignments require extensive data collection for multiple 2D and 3D NMR experiments on a single sample. With the small amount of CF present in the NMR sample (e.g. ∼70 nmoles of CF in a 1.6 mm rotor), 2D NCA, NCO and CC experiments can take several hours each, and 3D experiments such as NCACX may require days of data collection. Consequently, maintaining sample stability over a period of several days to weeks is crucial. Continuing sensitivity improvements will shorten these data collection times (26). Nevertheless, it is essential to be able to distinguish changes in samples due to deterioration from the changes due to intentional adjustment of the sample conditions, such as in the kinase-on and kinase-off states that we have examined in this study.

Functional and homogeneous samples of chemoreceptor signaling complexes are required for SSNMR studies investigating differences between signaling states. Microcrystalline samples often provide excellent resolution for SSNMR, but crystals reported for the chemoreceptor system include a mixture of native and non-native structures (16, 39). Functional complexes of nano disc-imbedded intact receptors have been studied, but these include a statistical mixture of receptors inserted in both directions, leading to sample inhomogeneity (40). In contrast, *in vitro* assembly of functional complexes of the Asp receptor CF with CheA and CheW yields homogeneous functional signaling arrays well-suited for SSNMR. These arrays, formed either through binding of the CF His tag to unilamellar lipid vesicles or by molecular crowding agents such as PEG8000, activate both the kinase CheA and methylation of the CF (18, 19). The arrays can be pelleted by ultracentrifugation to maximize the amount of sample packed in the SSNMR rotor while maintaining kinase and methylation activity.

Stable sample conditions for CF4Q assemblies (kinase-on) were identified through measuring the kinase activity of CheA, which is only activated in complexes, with an ATP consumption assay. Activity of the complexes was monitored over the course of 30 days, for samples kept in solution or as pellets. The latter was done by pelleting a series of aliquots and storing the supernatant and pellet separately at 4°C, then resuspending each pellet with its corresponding supernatant before measuring the kinase activity. Interestingly, vesicle assemblies of kinase-on complexes retain activity in solution for long periods of time, but after storage in the pelleted form, the resuspended assemblies lose most of their activity in three days (Fig 2A). This indicates that samples prepared with vesicle-mediated assembly are not suitable for long SSNMR experiments. In contrast, PEG8000-mediated kinase-on assemblies retain full kinase activity when stored in solution, and about 85% activity when stored as pellets for up to 30 days. Kinase-off (CF4E) PEG8000-mediated assemblies also maintain about 70% methylation activity over long storage periods as a pellet (Fig 2A). Methylation activity was determined by the ratio of methylated to unmethylated CF observed by gel electrophoresis, 4 hours after the addition of the methyltransferase, CheR, and the methyl group donor S-adenosyl methionine (SAM) (Fig S1). Notably, tests performed at higher storage temperatures (25°C and 12°C, data not shown) demonstrated an increasing loss of activity with increasing temperature.

**Figure 2.**
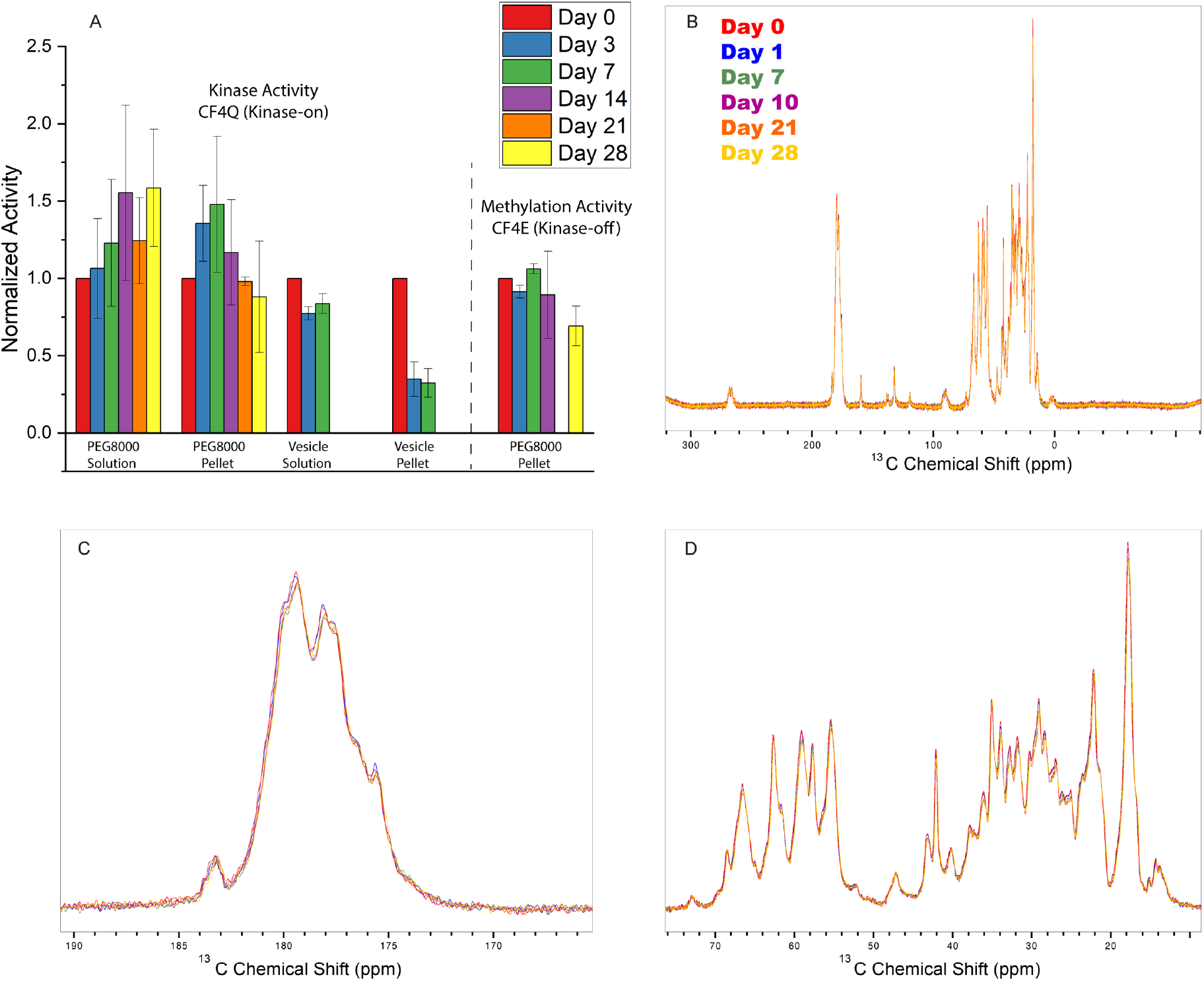
Sample stability monitored by activity assays and NMR spectra. (A) Kinase activity assays of CF4Q assemblies and methylation activity assays of CF4E assemblies were performed on mirror samples for both lipid vesicle and PEG8000-mediated assemblies stored in solution and as pellets. Monitoring of vesicle assemblies was ended after 7 days due to the loss of activity of the pellets. Data bars represent the mean of 3 replicate stability studies, with error bars showing ±SD and with Day 0 activity normalized to 1 for both assays. (B) 1D ^1^H-^13^C CP spectra collected throughout the NMR data collection period for a PEG-mediated CF4Q assembly in a 1.6 mm rotor. Expansion of (C) carbonyl region and (D) aliphatic region.

These pellet stability studies demonstrate that the resuspended pellets have comparable activity to the pre-centrifugation arrays, and PEG8000-mediated assembly produces arrays of both signaling states that retain function over a time frame suitable for extensive SSNMR data collection. Thus, all NMR samples were prepared using PEG8000-mediated assembly of U-^13^C,^15^N-CF, with unlabeled CheA and CheW, and then assessed using sedimentation assays (to measure the protein stoichiometry in the array) and either kinase assays (for kinase-on complexes assembled with CF4Q) or methylation assays (for kinase-off complexes assembled with CF4E). Moreover, the actual sample temperature was maintained at 4°C to ensure optimal stability throughout NMR data collection.

To further verify sample integrity during SSNMR experiments, 1D ^13^C CP spectra and short 2D CC spectra were recorded periodically between multidimensional experiments over the 30-day data collection period (Figure 2B-D and Fig S2). The lack of significant changes in spectral intensity, peak linewidth, or chemical shifts confirmed that the sample remained stable throughout data acquisition. Additionally, 2D NCA spectra of two separately prepared PEG8000-mediated kinase-on assemblies, one in a 1.6 mm rotor and the other in a 2.5 mm rotor, yielded nearly consistent results, demonstrating the reproducibility of the assembly method (Fig S3). It should be noted that the 2.5 mm rotor can hold over twice as much sample as the 1.6 mm rotor, leading to increase in signal intensity and number of signals. Only about 4 out of the 96 total peaks in the 1.6 mm NCA spectrum do not clearly overlap with peaks in the 2.5 mm spectrum. Thus, there may be very small differences in the assemblies, but the majority of peaks agree. These findings confirm that PEG8000-mediated assembly and 4°C temperature conditions enable long-term SSNMR studies of functional, pelleted signaling complexes, ensuring sufficient stability for multiple 2D and 3D NMR experiments.

### Dynamics-based Spectral Editing Yields Spectra of Region of Interest

The 57-residue protein interaction region (PIR) of the chemoreceptor interacts with both CheA and CheW, making it a key area for understanding the mechanism of chemoreceptor control of the kinase activity. Previous NMR and mass spectrometry studies from our lab predicted that the PIR is part of the rigid region of CF. This rigidity should allow for selective detection and assignment of this region using dipolar coupling-based SSNMR experiments.

A 2D NCA spectrum of uniformly labeled CF4Q (kinase-on) in PEG8000-mediated complexes contains 96 peaks (Fig 3B-C and Table 1). The same spectrum collected on CF4E (kinase-off) complexes detected 82 peaks (Fig 4B-C and Table 1). These results indicate that approximately 25-30% of the 310 total residues in CF are rigid, consistent with our prediction that a small fraction of CF is rigid in functional complexes. Amino acid type assignments were made for amino acids with unique chemical shifts in the NCA spectrum or unique C_β_ chemical shifts, using CC-DARR and NCACX experiments (Table S1). Table 1 compares the residue counts for glycine, alanine, threonine, serine and valine residues detected in CF4Q and CF4E spectra to their occurrences in the entire CF sequence and the PIR. The numbers of these amino acid resonances in the NMR spectra matches or exceeds their expected occurrence in the PIR, suggesting that these dipolar coupling-based experiments capture not only the PIR, but also an additional 20-30 amino acids within CF. Identification of unique dipeptides and tripeptides using connections given by NCO (Figure S4) and CANCO spectra have enabled us to assign about 40 of the 96 resonances in the CF4Q (kinase-on) NCA spectrum (Table S2). The assigned resonances localize within the PIR and part of the adjacent flexible bundle region, extending up to the glycine hinge on either side of the PIR (Figure 3A). This region will subsequently be referred to as the PIR-plus. The amino acid composition of the PIR-plus closely matches that of the detected residues in the NCA spectra (Table 1). Further data collection is needed to complete the assignment of these spectra and confirm whether only the PIR-plus region is detected in NMR spectra that require rigid-limit CN dipolar couplings.

**Figure 3.**
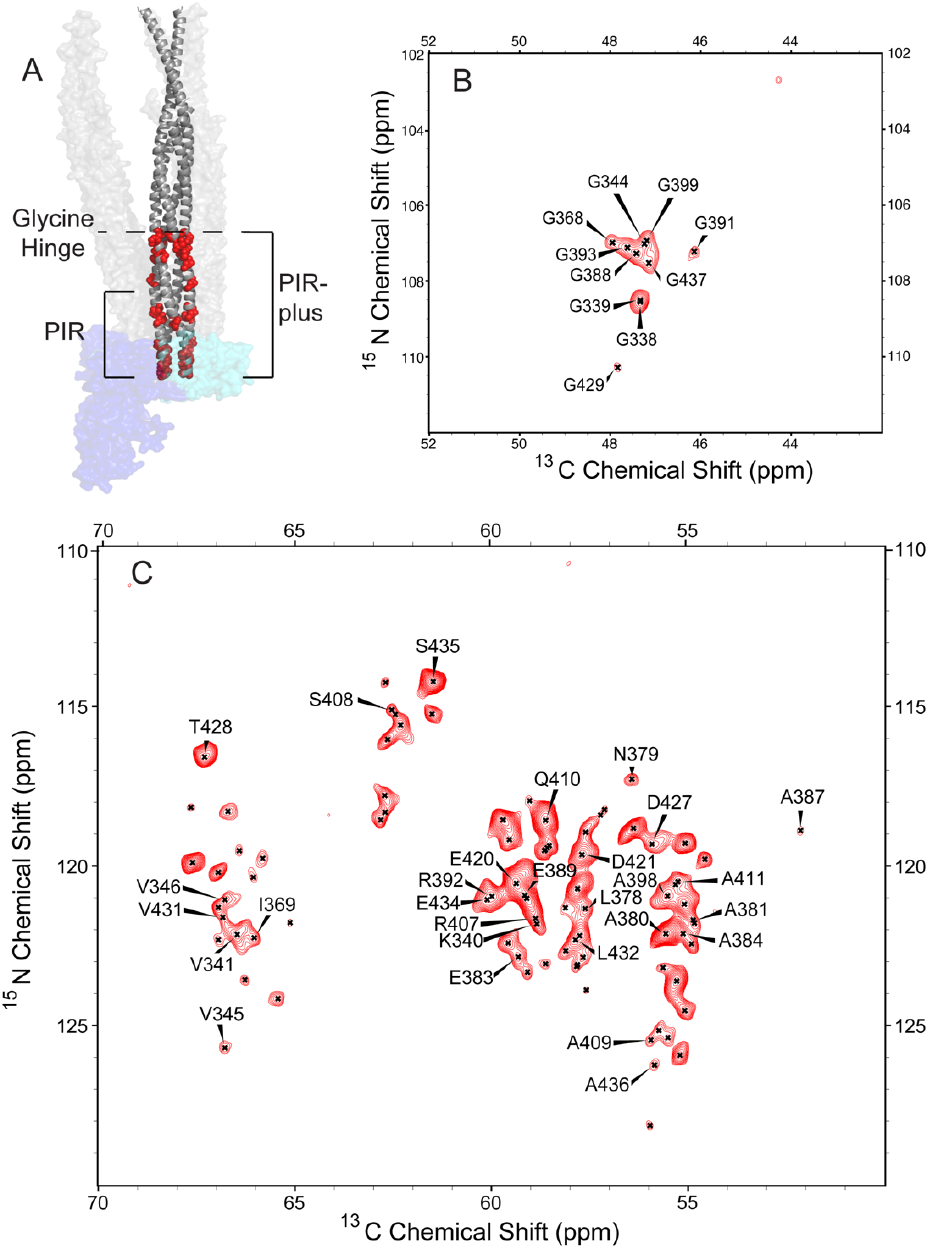
Assignments of CF4Q in PEG8000-mediated assemblies. (A) Dimer of CF (gray cartoon) in the context of a trimer-of-dimers (other dimers shown as transparent gray surface) and the other proteins of the complex (CheA, transparent blue surface and CheW, transparent cyan surface; PDB ID 6S1K, with 1QU7 used to extend to full length CF and residue numbers changed to aspartate chemoreceptor numbers). Assigned residues (backbone atoms shown as red spheres) were identified by a suite of 2D and 3D experiments. All assigned residues are between the glycine hinge (dashed line) and the tip of the PIR. (B) Glycine region and (C) main region of 2D NCA spectrum with assignments.

**Table 1:** Amino acid types identified in 2D NCA spectra^1^ compared to CF.

| Amino Acid | CF4Q NCA | CF4E NCA | Full CF | PIR<sup>2</sup> | PIR-plus<sup>3</sup> |
| --- | --- | --- | --- | --- | --- |
| Glycine | 10 | 8 | 17 | 5 | 10 |
| Alanine | 19 | 15 | 52 | 14 | 17 |
| Threonine | 3 | 3 | 21 | 1 | 3 |
| Serine | 9 | 8 | 31 | 3 | 9 |
| Valine | 12 | 11 | 24 | 5 | 13 |
| All Residues | 96 | 82 | 310 | 57 | 100 |

<sup>1</sup>The detection threshold was set to $7\sigma$ for both spectra for comparison.
<sup>2</sup>PIR defined as *E. coli* aspartate chemoreceptor residues 361-417
<sup>3</sup>PIR-plus defined as *E. coli* aspartate chemoreceptor residues 338-437

**Figure 4.**
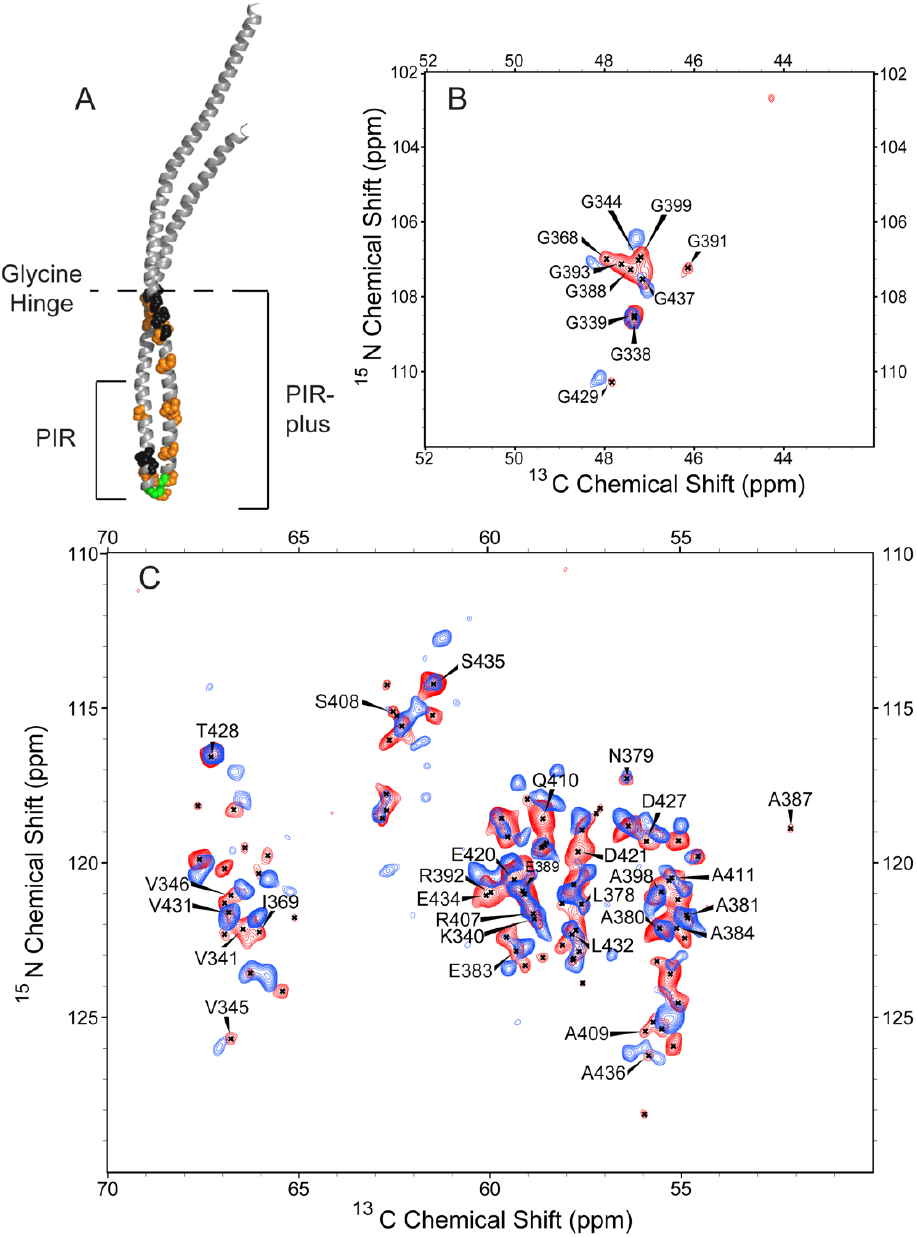
Comparison of NMR spectra of CF4Q and CF4E in complexes. (A) A monomer of CF with residues assigned in CF4Q shown as spheres. Assigned residues with no significant shifts between signaling states are shown in black, residues that have chemical shift changes between signaling states are orange, and residues that only have peaks in the kinase-on state are in green. (B-C) Overlay of 2D NCA spectra of CF4Q (red) and CF4E (blue) in PEG8000-mediated assemblies for (B) glycine region and (C) main region of 2D NCA, with assignments for CF4Q spectrum.

### NMR-Detected Region Reveals Signaling-Related Conformational Changes

Although the PIR is the most well-ordered region of CF observed by cryo-ET studies of these signaling complexes, at the limited resolution of the cryo-ET (∼8 Å at best) there are no detectable structural changes in this region for different signaling states. SSNMR, with its atomic level detection, provides a powerful approach to reveal subtle signaling-related changes in the protein-protein interfaces. Comparison of 2D NCA spectra of CF in kinase-on and kinase-off signaling complexes demonstrates that SSNMR detects significant changes (Fig 4B-C). Based on our current residue assignments for CF4Q, the following changes occurring between signaling states are highlighted in Figure 4A and Table S3. Out of the 96 peaks present in the CF4Q NCA spectrum, only about 10 peaks have little to no chemical shift changes between signaling states. These residues (black spheres in Figure 4A) are primarily located above the PIR near the glycine hinge. In contrast, residues that show notable chemical shift perturbations between states (orange spheres in Figure 4A) are widespread in the PIR-plus, suggesting that this region is experiencing conformational changes in addition to changes at the protein-protein interfaces. Finally, the NCA spectrum of CF4E (kinase-off) complexes contains fewer peaks than that of CF4Q complexes, which may indicate that a portion of the region being observed is more dynamic in the kinase-off state, causing the suppression of rigid-limit dipolar couplings. Three assigned residues in the turn between helices at the tip of the PIR are among the resonances missing in the spectrum of the kinase-off state (green spheres in Figure 4A), suggesting this turn is more dynamic in the off state. Further assignment of these spectra will allow us to identify which residues show chemical shift perturbations and which undergo changes in dynamics, providing deeper insight into the molecular mechanisms underlying chemoreceptor signaling.

## Discussion

The ability to study functional *in vitro* chemoreceptor signaling complexes is central to this work. Here we have demonstrated that pellets of PEG8000-mediated *in vitro* assemblies maintain kinase activity (kinase-on state) or methylation activity (kinase-off state) throughout the duration of SSNMR experiments. This assembly method also yields reproducible samples, based on the similarity of spectra of two separately prepared kinase-on assemblies. The agreement between NMR spectra interspersed throughout the long experiment times indicate that the sample integrity is maintained throughout the data collection under MAS conditions, resulting from a combination of the improvements in sample preparation and use of a low electric field NMR probe (28). In addition, the negligible kinase activity of PEG-mediated CF4E assemblies make these a better representative of a kinase-off state (vesicle-mediated assemblies often retain 50% kinase activity) (41). Moreover, PEG-mediated assemblies allow ∼50% more sample to be packed into the NMR rotor compared to vesicle-mediated assemblies (14).

It is interesting that lipid vesicle assemblies lose activity quickly in pelleted form, but not in solution. Cryo-EM images have revealed that unilamellar vesicles transform into multilamellar vesicles after several days of magic angle spinning (42). Perhaps such a transformation also occurs in the pellet samples we evaluated for activity, because the loss of accessibility of the kinase in multilamellar vesicles would lead to loss of measurable kinase activity in our assay. Such a transformation in the sample pellets, with or without MAS, would be problematic for our system, because the close separation between bilayers in multilamellar vesicles would very likely distort the 200 Å long CF complexes. Thus, pellets of vesicle-mediated assemblies are not suitable for long-duration SSNMR studies.

Here we have demonstrated that selective detection of the most rigid regions of a partially dynamic protein, combined with data collection at higher magnetic field, can yield simplified spectra to enable sequence-specific assignments of functionally important regions. Previous work in our lab predicted that about 80 residues of the CF of the chemoreceptor in functional complexes would be observed in NMR experiments that involve magnetization transfers mediated by ^13^C-^15^N dipolar couplings (15, 22). This was calculated based on estimates that ∼40% of its 310 residues are CP-observable (from the intensity ratio of unfrozen/frozen CP spectra), and that 60-65% of the CP-observable residues have rigid-limit dipolar coupling (from ^13^C-^15^N REDOR experiments). Indeed, the NCA spectrum of kinase-on CF4Q complexes displays good resolution, with resonances for ∼96 of the 310 residues of CF. Analysis of these spectra enable us to test our prediction that the observed rigid region includes the PIR, as this region exhibits significantly slower hydrogen exchange than the rest of the CF (23).

Amino acid type assignments of the 2D NCA spectrum of CF4Q as well as the sequence-specific assignment of 40 residues, using multiple 2D and 3D experiments, have demonstrated that dipolar coupling-based NMR spectra selectively detect the PIR-plus, consisting of the protein interaction region and adjacent segments extending to the glycine hinge. This region represents about a third of the residues present in the CF. This demonstrates that for dynamic proteins with a range of different motional timescales, dipolar coupling-based NMR experiments can be used to focus on regions with slower than millisecond motion to greatly simplify spectra of large proteins.

SSNMR of the PIR within functional complexes is a promising approach to reveal signaling-related changes that have not emerged from Cryo-ET. Cryo-ET studies of lysed cells and *in vitro* complexes have yielded structural models of chemoreceptor signaling complexes that are essential for understanding signaling mechanisms (16, 17, 43–45). While the PIR of CF is resolved in these studies, the resolution is insufficient to observe specific changes in its protein interfaces or dynamics between signaling states. Cryo-ET of *in vitro* assemblies of the CF of aspartate chemoreceptors with its associated proteins bound to lipid monolayers have led to 8.4 Å and 14.5 Å density maps for complexes with CF4Q and CF4E, respectively. The lower resolution of the CF4E complexes suggests these assemblies are less ordered than the CF4Q assemblies, and has made comparison of structures difficult (17). Cryo-ET of full length serine chemoreceptor complexes from lysed *E. coli* cells resulted in 20 Å and 24 Å density maps for CF4Q and CF4E, respectively, that showed complexes with 4E having tighter packing of trimers-of-dimers below a bent glycine hinge, and more dynamic methylation helices compared to 4Q (46). The PIR in these density maps are very similar. In contrast, SSNMR spectra of the CF in kinase-on and kinase-off signaling states show many chemical shift changes. This indicates multiple small changes in conformation and protein interactions are occurring in this region that are too subtle to be resolved by cryo-ET. Assigning these spectra and combining them with current models of the complexes will identify residues involved in changes that propagate the signal to control the kinase.

In addition to chemical shift changes, NMR spectra of the rigid region of CF in kinase-off complexes show a reduced number of peaks that likely indicates increased mobility. Our lab has shown that CF in kinase-on complexes is more rigid than in kinase-off complexes and hypothesized that reduced dynamics are important for kinase activation (14, 15, 23, 41). We previously identified regions of CF with rapid dynamics as those that are observed in scalar coupling-based INEPT NMR spectra and exhibit very rapid hydrogen exchange. Methylation helix 1 (MH1) and the C-terminal tail of CF exhibits such dynamics in both signaling states; methylation helix 2 (MH2) exhibits such dynamics in kinase-off (CF4E) signaling complexes, accounting for some of the difference in the overall dynamics of CF between methylation states. Here, we show that the PIR-plus, the membrane-distal tip region of the CF, loses about 12 rigid residues in the kinase-off compared to kinase-on signaling complexes. Residues near the glycine hinge, which appears to be the end of the rigid PIR-plus region, are detected in NMR spectra of both signaling states. This suggests that it is not these edge residues of the rigid region that are becoming dynamic. The 12 peaks lost in spectrum of the kinase-off state compared to the kinase-on state include 4 Ala and 2 Gly, similar to the composition of the turn between helices at the receptor tip (384-394, containing 3 Ala and 3 Gly). Furthermore, 3 of the residues that have been assigned in this region in CF4Q (A387, G388 and G391) appear to be missing in CF4E. This may indicate that the membrane-distal tip of the chemoreceptor undergoes changes in dynamics between signaling states, with increased dynamics and weaker protein-protein interactions in the kinase-off state. This is in line with our hypothesis that the tip of the chemoreceptor is more dynamic in the kinase-off state, which weakens the protein-protein interactions and allows the kinase to relax into the off state. Further NMR experiments, including applying selective isotope labeling and spectral editing approaches, will be pursued to facilitate the resonance peak assignments to test this hypothesis and to determine what changes occur in the important contacts in signaling complexes.

## Conclusion

This study demonstrates the promise of high-field (900 MHz) SSNMR as a powerful tool for mechanistic studies of multiprotein complexes. Such studies are critical because biological function in cells occurs within large assemblies of multiple proteins. In the case of bacterial chemotaxis receptor signaling complexes, Cryo-ET has provided valuable structural models but has not been able to resolve structural changes at the receptor/kinase interfaces that regulate kinase activity. We have developed optimized sample preparation methods that support extended SSNMR data collection on active, native-like chemoreceptor complexes, utilizing a low electric field probe design to reduce sample damage during data collection. Because the receptor CF exhibits significant dynamics in functional complexes, NMR methods for selective detection of the most rigid region greatly simplify the spectrum and have enabled assignment of ∼40% of the observed resonances. These assignments demonstrate that the rigid region includes the receptor protein interaction region that is integral to the control of the kinase in this system. Significant changes in the NMR spectra between different signaling states indicate that NMR is a fruitful approach for mapping subtle structural changes in the receptor and its contacts, to complement existing studies and reveal the signaling mechanism. Such an approach could be broadly applied to other large proteins and their complexes, enabling selective investigation of mechanistically important regions and functionally relevant dynamics.

## Author Contributions

J.J.A prepared samples, performed sample stability studies, analyzed the NMR data, and wrote the manuscript. S.W. performed NMR data collection, processed NMR data, and contributed to writing the manuscript. L.K.T. supervised the research, acquired funding, and edited the manuscript. C.M.R. supervised the research, acquired funding, installed the spectrometers and probes, and edited the manuscript. All authors contributed to the final version of the manuscript.

## Declaration of Interests

CMR is a founder of Resynant, Inc. and a member of its scientific advisory board.

## Acknowledgements

This work was supported by NIH Grant R01-GM120195 (to LKT). JJA was partially supported by National Research Service Award T32 GM139789 from the National Institutes of Health. This study made use of the National Magnetic Resonance Facility at Madison (NMRFAM), an NIH Biomedical Technology Development and Dissemination Center (P41GM136463). Helium recovery equipment, computers, and infrastructure for data archive were funded by the University of Wisconsin-Madison, NIH (P41GM136463, R24GM141526), and NSF (1946970).

