## Supplemental Information for "Selective Detection of a Key Region in Chemotaxis Signaling Protein Complexes by Solid-State NMR"

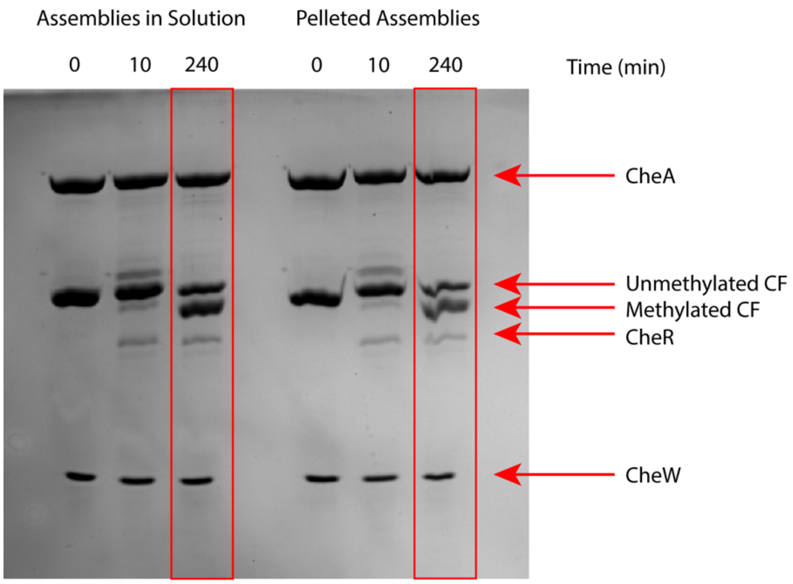


**Figure S1**. Example of methylation activity assay gel. Methylation activity assay of CF4E (kinase-off) PEG8000-mediated assemblies stored at 4°C for 3 days in solution (left) and as a pellet (right). Pellets were resuspended with their supernatant prior to the assay. Gel samples were taken before initiation of the assay, 10 minutes after CheR and SAM were added to begin methylation, and 240 minutes later. Methylation activity was calculated based on the ratio of the unmethylated and methylated bands in the 240 minutes columns. Arrows indicate which bands correspond to the proteins of the complexes as well as the methyltransferase, CheR.

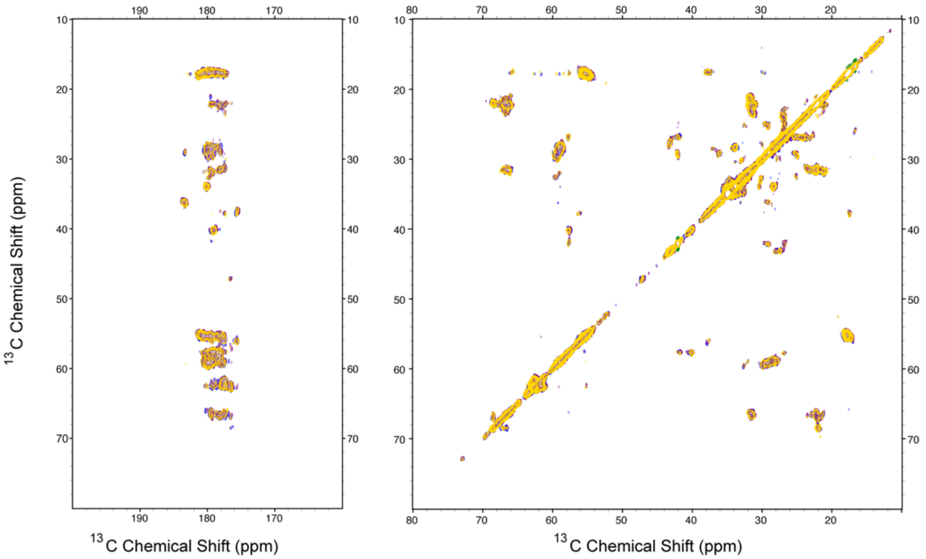


**Figure S2**. 2D CC experiments collected between 3D experiments to monitor sample integrity, carbonyl region in left panel and aliphatic region in right panel. The blue spectrum was collected on the 1st day of the data collection period, the purple spectrum was collected on the 10th day, and the yellow spectrum was collected on the 21st day.

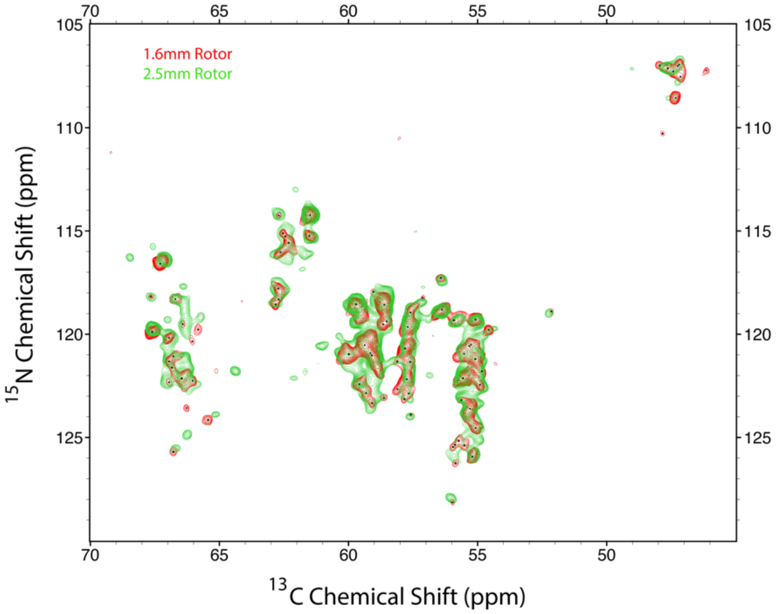


**Figure S3**. 2D NCA spectra of separately assembled CF4Q PEG8000-mediated assemblies. Both spectra were collected at 900MHz with 20kHz magic angle spinning. The red spectrum was collected on ~70 nmol CF in complexes, using a Blackfox 1.6mm HXY probe; the green spectrum was collected on ~170 nmol CF in complexes, using a Blackfox 2.5mm HXY probe.

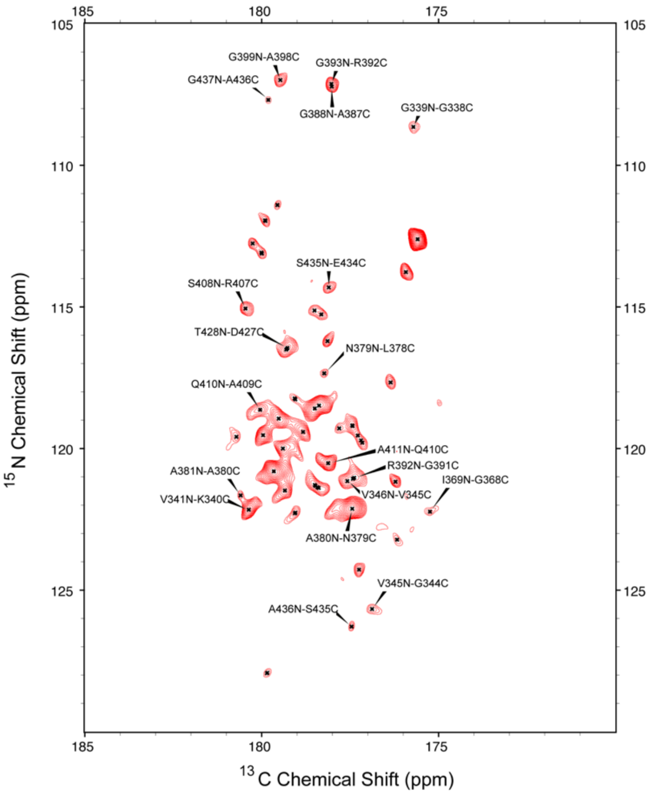


**Figure S4**. 2D NCO spectrum of CF4Q PEG8000 assembly with assignments.

**Table S1**. List of CF4Q assignments of resonances in intra-residue experiments.

| Residue | NCA<br>N(ppm), C(ppm) | CC DARR<br>C(ppm), C(ppm) | NCACX<br>N(ppm), C(ppm), C(ppm) |
|---|---|---|---|
| G338 | 108.553, 47.337 | | 108.577, 47.413, 47.486 N-CA-CA<br>108.682, 47.416, 175.715 N-CA-CO |
| G339 | 108.512, 47.338 | | |
| K340 | 121.813, 58.845 | 58.891, 31.793 CA-CB<br>58.952, 28.295 CA-CD<br>31.813, 58.857 CB-CA<br>31.811, 21.228 CB-CG<br>28.309, 58.907 CD-CA<br>28.174, 43.368 CD-CE<br>43.367, 28.171 CE-CD<br>21.231, 31.798 CG-CB | 121.836, 58.937, 58.907 N-CA-CA<br>121.875, 58.879, 31.87 N-CA-CB<br>121.836, 58.915, 180.379 N-CA-CO |
| V341 | 122.154, 66.467 | 66.555, 31.449 CA-CB<br>66.502, 22.107 CA-CG<br>31.35, 66.496 CB-CA<br>22.048, 66.395 CG-CA | 122.041, 66.465, 66.443 N-CA-CA<br>122.506, 66.699, 31.514 N-CA-CB |
| G344 | 107.021, 47.241 | | 107.125, 47.199, 47.172 N-CA-CA<br>107.275, 47.214, 176.871 N-CA-Co |
| V345 | 125.702, 66.787 | 66.779, 177.608 CA-CO<br>66.767, 23.319 CA-CG<br>23.537, 66.726 CA-CG<br>31.457, 66.767 CB-CA | |
| V346 | 121.066, 66.787 | | 121.059, 66.744, 66.685 N-CA-CA<br>121.153, 66.891, 31.917 N-CA-CB |
| G368 | 106.988, 47.951 | 47.929, 175.2 CA-CO | |
| I369 | 122.259, 66.036 | 66.058, 17.792 CA-CD<br>66.068, 179.452 CA-CO<br>37.763, 66.079 CB-CA<br>17.864, 66.06 CD-CA | |
| L378 | 121.342, 57.613 | 57.725, 41.78 CA-CB<br>57.658, 178.221 CA-CO<br>41.866, 57.65 CB-CA<br>41.836, 178.233 CB-CO | |
| N379 | 117.274, 56.464 | 56.451, 38.079 CA-CB<br>56.447, 177.523 CA-CO<br>38.123, 56.493 CB-CA<br>38.077, 177.558 CB-CO | 117.572, 56.464, 56.455 N-CA-CA<br>117.388, 56.474, 38.388 N-CA-CB |
| A380 | 122.131, 55.57 | | 122.142, 55.542, 55.53 N-CA-CA<br>122.225, 55.515, 17.836 N-CA-CB<br>122.115, 55.561, 180.572 N-CA-CO |
| A381 | 121.7, 54.872 | | |

| | | | |
|---|---|---|---|
| E383 | 122.854, 59.323 | 59.246, 29.233 CA-CB<br>59.324, 36.348 CA-CG<br>59.319, 178.364 CA-CO<br>29.255, 59.256 CB-CA<br>29.231, 36.167 CB-CG<br>36.453, 59.448 CG-CA<br>36.247, 29.163 CG-CB | 123.043, 59.32, 59.29 N-CA-CA<br>123.084, 59.244, 29.413 N-CA-CB |
| A384 | 122.12, 55.123 | 55.181, 17.823 CA-CB<br>17.906, 55.132 CB-CA | 122.12, 55.016, 54.989 N-CA-CA<br>122.12, 55.084, 18.332 N-CA-CB |
| A387 | 118.898, 52.139 | 52.321, 19.082 CA-CB<br>52.299, 178.192 CA-CO<br>19.019, 52.162 CB-CA | 118.758, 52.286, 52.286 N-CA-CA<br>118.953, 52.316, 19.247 N-CA-CB |
| G388 | 107.277, 47.424 | | 107.209, 47.429, 177.827 N-CA-CO |
| E389 | 120.915, 59.157 | | 120.762, 59.098, 28.388 N-CA-CB<br>120.745, 59.049, 176.575 N-CA-CO |
| G391 | 107.227, 46.14 | 46.139, 177.309 CA-CO | 107.422, 46.152, 46.102 N-CA-CA |
| R392 | 120.963, 59.985 | 59.92, 29.856 CA-CB<br>29.837, 59.919 CB-CA<br>29.741, 41.963 CB-CD | 120.971, 60.069, 60.039 N-CA-CA<br>121.069, 60.028, 178.069 N-CA-CO |
| G393 | 107.123, 47.618 | | 107.178, 47.583, 47.55 N-CA-CA<br>107.238, 47.614, 175.902 N-CA-CO |
| A398 | 120.942, 55.526 | | 120.996, 55.505, 55.534 N-CA-CA<br>120.996, 55.12, 17.667 N-CA-CB<br>120.996, 55.41, 179.605 N-CA-CO |
| G399 | 106.937, 47.194 | | |
| R407 | 121.645, 58.883 | 58.811, 31.734 CA-CB<br>58.736, 180.429 CA-CO<br>31.854, 180.445 CB-CO | |
| S408 | 115.11, 62.539 | | 115.1, 62.634, 62.642 N-CA-CA |
| A409 | 125.458, 55.946 | | 125.64, 55.891, 55.901 N-CA-CA<br>125.429, 55.881, 180.021 N-CA-CO |
| Q410 | 118.577, 58.621 | 58.607, 178.31 CA-CO<br>58.614, 33.79 CA-CG<br>34.023, 58.579 CG-CA | 118.672, 58.662, 58.65 N-CA-CA<br>118.561, 58.691, 28.926 N-CA-CB<br>118.589, 58.566, 178.217 N-CA-CO |
| A411 | 120.492, 55.257 | | 120.5, 55.263, 55.276 N-CA-CA<br>120.5, 55.064, 18.054 N-CA-CB |
| E420 | 120.544, 59.37 | 59.348, 32.14 CA-CB<br>59.371, 36.396 CA-CG<br>59.38, 180.035 CA-CO<br>32.205, 59.372 CB-CA | 120.651, 59.408, 59.405 N-CA-CA<br>120.477, 59.414, 29.29 N-CA-CB<br>120.709, 59.474, 180.28 N-CA-CO |
| D421 | 119.646, 57.7 | 57.706, 40.291 CA-CB<br>57.711, 179.52 CA-CO<br>40.247, 57.677 CB-CA | 119.559, 57.714, 57.723 N-CA-CA<br>120.054, 57.697, 40.599 N-CA-CB<br>119.649, 57.604, 179.333 N-CA-CO |

| | | | |
|---|---|---|---|
| D427 | 119.321, 55.919 | 55.909, 37.603 CA-CB<br>37.513, 55.948 CB-CA | 119.291, 55.987, 55.999 N-CA-CA<br>119.123, 56.042, 37.6 N-CA-CB<br>119.377, 55.884, 179.235 N-CA-CO |
| T428 | 116.444, 67.158 | 67.108, 68.581 CA-CB<br>68.6, 67.133 CB-CA | 116.583, 67.283, 67.317 N-CA-CA<br>116.612, 67.361, 68.651 N-CA-CB |
| G429 | 110.289, 47.837 | | |
| V431 | 121.613, 66.837 | 66.759, 178.27 CA-CO | |
| L432 | 122.313, 57.862 | 57.691, 41.876 CA-CB<br>41.866, 57.707 CB-CA<br>26.739, 57.886 CD-CA | 122.033, 57.741, 57.784 N-CA-CA<br>121.908, 57.722, 41.878 N-CA-CB |
| E434 | 121.064, 60.108 | 60.144, 177.997 CA-CO | |
| S435 | 114.232, 61.482 | 61.517, 62.664 CA-CB<br>61.513, 177.527 CA-CO<br>62.665, 61.482 CB-CA | 114.352, 61.563, 61.569 N-CA-CA<br>114.332, 61.574, 62.804 N-CA-CB<br>114.298, 61.545, 177.436 N-CA-CO |
| A436 | 126.244, 55.854 | | 126.436, 56.283, 56.116 N-CA-CA<br>126.38, 55.943, 18.173 N-CA-CB<br>126.237, 56.042, 180.574 N-CA-CO |
| G437 | 107.53, 47.145 | | |

**Table S2**. List of CF4Q assignments of resonances in inter-residue experiments.

| Residues | NCO<br>N(ppm), C(ppm) | NCOCX<br>N(ppm), C(ppm), C(ppm) | CANCO<br>C(ppm), N(ppm), C(ppm) |
|---|---|---|---|
| G339N-G338C | 108.64, 175.709 | 108.746, 175.72, 175.839 | |
| V341N-K340C | 122.16, 180.36 | 122.139, 180.43, 180.406 | |
| V345N-G344C | 125.669, 176.88 | 125.483, 176.904, 176.866<br>125.83, 177.042, 47.176 | |
| V346N-V345C | 121.142, 177.581 | | |
| I369N-G368C | 122.229, 175.246 | | |
| N379N-L378C | 117.344, 178.226 | | |
| A380N-N379C | 122.118, 177.441 | | 55.612, 122.175, 177.643 |
| A381N-A380C | 121.65, 180.597 | | |
| A384N-E383C | | | 55.126, 122.106, 178.35 |
| G388N-A387C | 107.221, 178.023 | | |
| E389N-G388C | | | 59.387, 120.913, 177.822 |
| R392N-G391C | 121.028, 177.381 | | |
| G393N-R392C | 107.124, 178.028 | | |
| G399N-A398C | 106.991, 179.474 | | 47.279, 107.291, 179.568 |
| S408N-R407C | 115.054, 180.456 | 115.183, 180.535, 180.469 | |
| Q410N-A409C | 118.633, 180.041 | | |
| A411N-Q410C | 120.515, 178.116 | 120.5, 178.163, 178.18<br>120.521, 178.205, 58.639 | |
| D421N-E420C | | | 57.647, 119.509, 180.232 |
| L432N-V431 | | | 57.815, 122.427, 178.274 |
| T428N-D427C | 116.442, 179.266 | 116.512, 179.233, 179.213 | |
| S435N-E434C | 114.313, 178.102 | | |
| A436N-S435C | 126.28, 177.457 | | |
| G437N-A436C | 107.688, 179.813 | | |

**Table S3**. List of what residues assigned in CF4Q stay the same or change compared to CF4E.

| | |
|---|---|
| No chemical shift change[1] | G338, G339, L378, N379, A380, A381, T428 |
| Chemical shift changes[2] | K340, V341, G344, V345, V346, G368, I369, E383, A384, E389, R392, G393, A398, G399, N403, A412, K413, E420, D421, D427, G429, V431, L432, G437 |
| Peak only found in CF4Q[3] | A387, G388, G391 |

[1] Peak overlap in CF4Q and CF4E 2D NCA spectra.

[2] Peaks shift in CF4Q versus CF4E 2D NCA spectra.

[3] Peaks are only seen in CF4Q 2D NCA spectrum and are absent in CF4E spectrum.
